# Binary node clustering via contrastive learning for haplotype phasing in de novo genome assembly

**DOI:** 10.64898/2026.07.17.739091

**Authors:** Martin Schmitz, Leon Rauschning, Kenji Kawaguchi, Mile Sikic

## Abstract

Accurate haplotype phasing is essential for high-quality genome assembly, yet de novo phasing of complex genomes without parental data remains challenging. We formulate haplotype phasing as a node clustering problem with overlapping clusters on augmented unitig graphs, where nodes represent contiguous, non-branching DNA sequence fragments and two edge types can encode sequence overlap or Hi-C proximity information. We introduce a contrastive learning framework with a custom objective function and train a graph-transformer–based model, termed `grapHiC,` to phase paternal, maternal, and homozygous unitig nodes.

`grapHiC`; is the first machine-learning–based method to perform reference-free haplotype phasing and the first approach to directly phase raw unitig graphs without prior simplification. We show that `grapHiC`accurately clusters nodes on human-genome-scale graphs and that its predictions can effectively guide phased de novo genome assembly, producing human assemblies with contiguity and phasing quality comparable to the state of the art when integrated with the DipGNNome assembler.

## 1. Introduction

Genome assembly, the reconstruction of an organism’s genomic sequence from a collection of error-prone sequencing reads, is a fundamental algorithmic challenge in computational biology. Recent advances have seen machine learning techniques integrated into multiple stages of the genome assembly pipeline (Schmitz et al., 2026; Vrček et al., 2025; Luo et al., 2024; Battistella et al., 2025; Xue et al., 2022; Ke & Vikalo, 2020a;b). Many organisms possess multiple copies of their genome, called haplotypes, which may differ due to variants inherited from different parents. Most animals, including humans, are diploid, carrying two copies of each autosomal chromosome. Separating these haplotypes, a process known as *phasing*, is a critical component of many genomics workflows.

Accurately untangling haplotypes during genome assembly remains challenging, particularly for complex genomes, despite rapid progress in sequencing technologies and assembly algorithms. The most straightforward approach to haplotype separation is trio binning, which leverages parental genomes to assign variants to their parental origin. However, this approach requires additional sequencing and is infeasible for wild, historical, or otherwise undocumented samples where parental data cannot be obtained.

Chromatin conformation capture sequencing (Hi-C) measures the frequency of physical contacts between pairs of genomic loci, reflecting their spatial proximity within the nucleus. This long-range linkage information can be exploited for haplotype phasing, as loci on the same haplotype tend to interact more frequently than loci on different haplotypes. Hi-C sequencing does not require additional samples and has become increasingly affordable and routine, but the resulting data are noisy and subject to various technical and biological biases.

Recent work has demonstrated the potential of graph neural networks (GNNs) for *de novo* genome assembly. Vrček et al. (2025) introduced a GNN-based framework for haploid assembly, while Schmitz et al. (2026) extended this paradigm to diploid genomes using trio binning. These approaches highlight the suitability of graph-based deep learning for reasoning over large and complex assembly graphs. However, learning-based haplotype phasing methods without parental data remain unexplored.

To address this gap, we introduce `grapHiC` a graph-transformer-based framework for reference-free haplotype phasing that operates directly on unsimplified unitig graphs and jointly models sequence overlap and Hi-C information. To our knowledge, `grapHiC` is the first deep-learning–based method to perform reference-free haplotype phasing. It is also the first method to perform the step of phasing directly on unitig graphs.

Our code, trained model, and dataset are available at https://github.com/lbcb-sci/grapHiC.</

## 2. Related Work

In recent years, several approaches have leveraged neural networks for genome assembly tasks. For *de novo* assembly, GNNome (Vrček et al., 2025) performs layout on an Overlap–Layout–Consensus (OLC) graph using a Graph Convolutional Network (GCN), while GTasm (Luo et al., 2024) utilizes a graph transformer. Šimunović et al. (Šimunović et al., 2025) focus similarly on the layout of De Bruijn graphs. However, these methods assemble multiple haplotypes into a single consensus. DipGNNome (Schmitz et al., 2026) is the first machine learning-based *de novo* assembler to create phased assemblies, though it requires parental information to perform trio binning.

Several deep-learning methods have been proposed for haplotype phasing in a reference-based setting. GAEseq (Ke & Vikalo, 2020b) and CAECseq (Ke & Vikalo, 2020a) utilize autoencoders for read clustering, while NeurHap (Xue et al., 2022) improves performance by framing phasing as a graph coloring problem. More recently, `ralphi` (Battistella et al., 2025) uses deep reinforcement learning to optimize the maximum fragment cut (MFC). All of these approaches rely on aligning reads or fragments to a reference genome, producing comparatively clean graphs. Reference-derived graphs differ fundamentally from *de novo* unitig graphs, which are substantially noisier and contain many spurious connections. As a result, these methods are not directly applicable to reference-free assembly graphs.

Alignment-based approaches can also introduce reference bias, particularly in repetitive or structurally variable regions, which propagates into downstream analyses and is difficult to correct. Several recent phasing tools (Lorig-Roach et al., 2024; Ouchi et al., 2023; Zhang et al., 2024; Zhang, 2018; Cheng et al., 2021; Antipov et al., 2024) incorporate Hi-C data and may optionally leverage parental sequencing. However, these methods typically rely on strong simplifying assumptions about the input assembly graph, such as bubble-chain or otherwise highly simplified representations, or require substantial manual intervention. Consequently, phasing is performed only after partial graph resolution, potentially discarding informative graph structure.

Current state-of-the-art assemblers such as `hifiasm`(Cheng et al., 2021) and `Verkko`(Antipov et al., 2024) successfully integrate phasing into the assembly process without relying on machine learning. Both tools perform phasing only after simplifying the assembly graph into constrained structures, effectively functioning similarly to the phasing tools described above. In contrast, we investigate haplotype phasing directly on unsimplified unitig graphs, allowing this information to be used during the path tracing and graph simplification steps of genome assembly, and avoiding the misjoined contigs mixing sequence from different haplotypes.

We frame phasing as a node clustering problem. Traditional clustering approaches typically rely on the homophily assumption. This means nodes with strong connections are more likely to belong to the same cluster. Spectral clustering (Ng et al., 2001) uses Eigenvectors of the graph Laplacian for clustering, while the Louvain algorithm (Blondel et al., 2008) detects communities by iteratively maximizing modularity. Label propagation methods (Zhu & Ghahramani, 2002) iteratively diffuse known labels across the graph under the assumption that adjacent nodes tend to share similar properties. In cases where clusters are defined not by direct connections but by structural roles or more complex interaction patterns, learned deep representations can offer a benefit. On these representations, simple clustering algorithms such as k-nearest neighbor (KNN) clustering (Peterson, 2009; Keller et al., 1985) can be used, enabling the detection of higher-order patterns beyond simple connectivity.

In assembly graphs where clusters are defined by haplotypes, the homophily assumption does not characterize them well enough. Instead, we use a representation learning strategy. Representation learning approaches for graph clustering can fall into different categories such as reconstructive methods, often based on autoencoders (e.g. (Kipf & Welling, 2016)), (generative) adversarial methods (Mukherjee et al., 2019), and contrastive learning approaches (Watteau et al., 2024). Each of these strategies seeks to map nodes to a lower-dimensional embedding space where clustering is easier, but approaches this in different ways. Foundational work in establishing contrastive learning was published in (Chopra et al., 2005; Hadsell et al., 2006). Early graph-specific contrastive methods include the Linear Relational Encoding (LRE) strategy (Paccanaro & Hinton, 2000) and TransE (Bordes et al., 2013), applied to knowledge graph completion.

Contrastive objectives are typically label-free and effective at learning invariant representations through data augmentations and multiple views of the same data. InfoNCE (Oord et al., 2018) is among the most widely used label-free contrastive loss functions. Supervised Contrastive Learning (SupCon) (Khosla et al., 2020) extends this framework by incorporating label information: while negative pairs are often constructed implicitly via data augmentation, available class labels can also be used to define positive and negative relationships.

## 3. Overview

Overlap graphs are directed graphs in which nodes represent DNA sequences. A directed edge is added from one node to another if their sequences overlap, that is, if the suffix of one sequence matches the prefix of the other.

Unitig graphs are a compressed but otherwise unsimplified representation of raw overlap graphs. A read overlap graph can be transformed into a **unitig graph** in two steps. First, transitive edges are removed.

Second, all maximal non-branching paths are collapsed into single nodes, called *unitigs*. By operating directly on the unitig graph, before any additional simplifications or reductions, we aim to avoid errors that could arise from misjoining fragments belonging to different haplotypes. Unitig graphs can be large and complex. To make them amenable to graph machine learning approaches, we choose a linear-time graph transformer architecture capable of integrating global information and augment the graph with additional Hi-C information. Hi-C connections are added between nodes, with the edge weight indicating the normalized strength of the Hi-C signal mapping to both incident unitigs.

We model the unitig graphs as **heterogeneous graphs** with a single node type (each node represents a unitig) and two distinct edge types: **overlap edges** represent sequence overlaps, and **Hi-C connection edges** represent long-range chromatin contacts. On these heterogeneous graphs, we formulate haplotype phasing as a binary node clustering problem with overlapping clusters, where each cluster corresponds to one parent’s haplotype. In homozygous regions, where the genomic sequence inherited from both parents is identical, a node may be assigned to both haplotypes, resulting in overlap between the two clusters.

Framing the task as a clustering problem is therefore essential. Standard supervised classification losses fail to accommodate the permutation invariance inherent to haplotypes, where class labels are interchangeable, and cannot explicitly model the presence of homozygous nodes. Without parental data, haplotypes are interchangeable. For two variants on the same chromosome, it is possible to infer whether they were inherited from the same or different parents, but not the parental origin itself. For variants on different chromosomes, Hi-C-based connections cannot be used for phasing, as they do not originate from the same DNA molecule.

This means that any valid haplotype assignment remains valid after a chromosome-level swap of maternal and paternal labels. To accommodate this symmetry, we introduce a custom objective inspired by pairwise contrastive losses that explicitly models both haplotype label invariance and homozygosity. We refer to this objective as the supervised binary pair loss (SBP-Loss). Figure 1 illustrates the overall training and inference pipeline of `grapHiC`.

**Figure 1.**
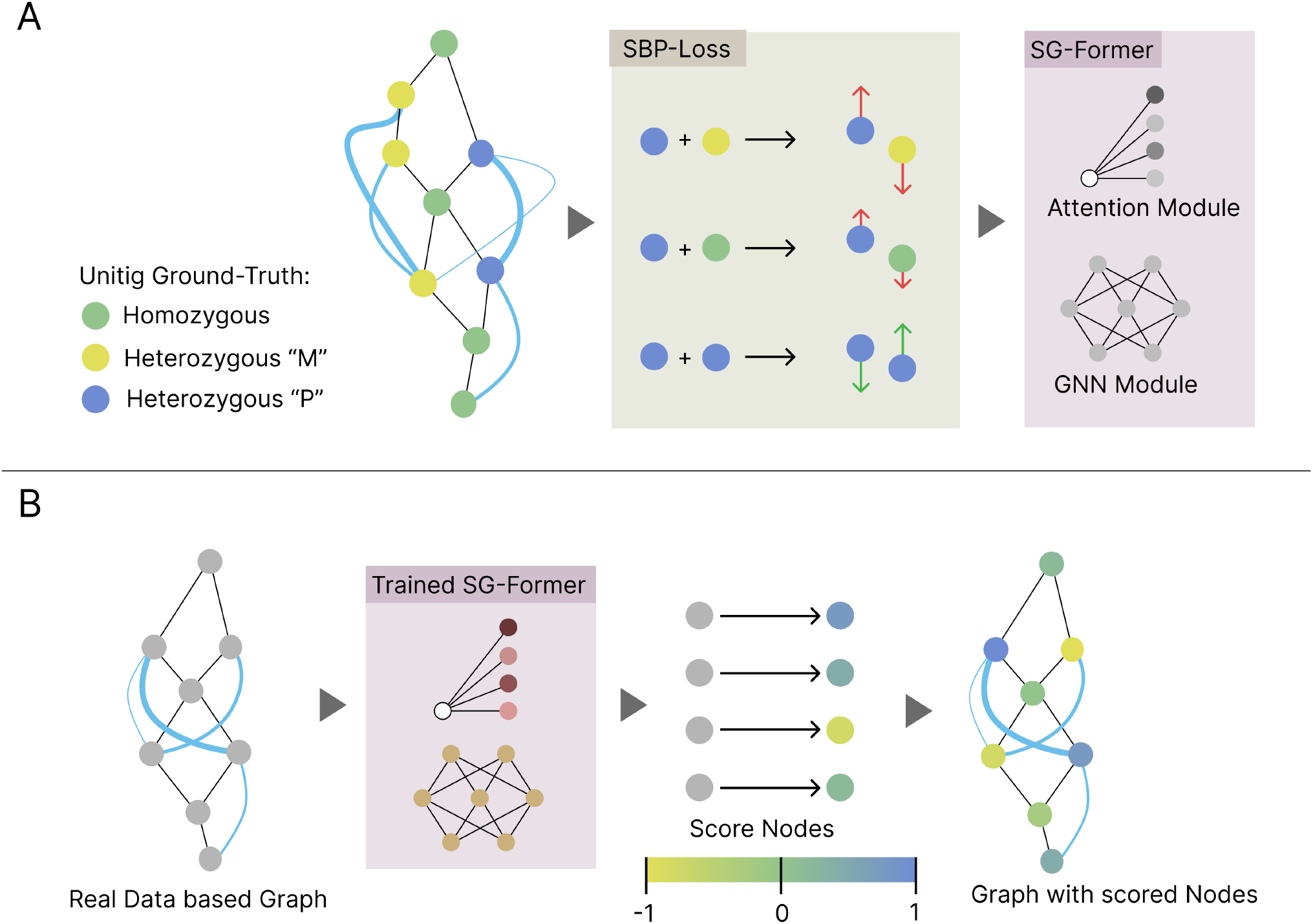
**A:** A training dataset of graphs is used to train an SG-Former (Wu et al., 2024) using SBP-Loss (proposed here) as training objective. **B:** The trained model gets an input graph, and outputs node scores, representing hapolotype allocations, and homozygousity probability.

In summary, our contributions are threefold:

1. we develop a data processing pipeline that constructs Hi-C–augmented unitig graphs as heterogeneous graphs suitable for machine learning workflows;
2. we introduce the first learning-based framework for reference-free haplotype phasing. It is also the first framework to perform phasing directly on unitig graphs; and
3. we propose a novel objective function tailored to binary (fuzzy) clustering, which outperforms standard contrastive losses in clustering accuracy by reflecting the symmetries and invariances of this problem and enables downstream assemblers to achieve state-of-the-art results in whole-genome phased *de novo* assembly.

## 4. Supervised Binary Pair Loss

We build upon the classic pair loss used in contrastive learning (Chopra et al., 2005; Hadsell et al., 2006), which aims to minimize distances between similar samples while enforcing a margin between dissimilar ones:

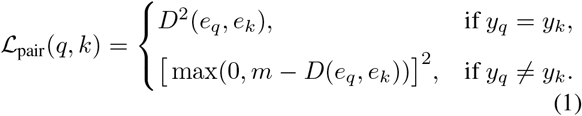

where *q* and *k* represent query and key samples, *eq, ek* are the corresponding embeddings, *yq, yk* are the class labels, *D* is the Euclidean distance, and *m* is a margin parameter.

During training, we have ground-truth labels for all nodes, distinguishing our haplotype phasing scenario from typical unsupervised clustering problems. Each node is assigned one of three labels: exclusively belonging to haplotype A (*y* = ***−***1), exclusively belonging to haplotype B (*y* = 1), or homozygous (being shared by both haplotypes) (*y* = 0). These labels can be leveraged in our objective function during training.

Given the binary nature of our clustering task, we employ a one-dimensional embedding space where the model outputs scalar predictions *p*∈ [*−*1; 1] for each node. Our goal is to push nodes from different haplotypes toward opposite extremes (*−* 1 and 1) while centering homozygous nodes around 0. For our one-dimensional case, the Euclidean distance reduces to the absolute difference *D*(*e*_*q*_, *ek*) = (*p*_*q*_ −*pk*).

We set the margin *m* = *|y_q_* − *yk*| ∈ {0, 1, 2}, which naturally encodes the desired distances.| When comparing nodes with identical labels (*y*_*q*_ = *y*_*k*_), we have *m* = 0, pushing the predicted labels towards exact agreement. For a pair consisting of a homozygous node (*y* = 0) and a heterozygous node exclusive to one haplotype (*y* = *±*1), we set *m* = 1. When comparing nodes from opposing haplotypes (*y* =***−*** 1 vs *y* = 1), we choose *m* = 2, requiring maximal separation.

By introducing an indicator variable *s*_*qk*_ to capture the relationship between labels, we can derive a unified formulation:

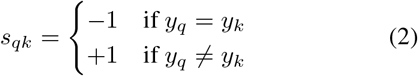

This allows us to express our *Supervised Binary Pair (SBP) Loss* as:

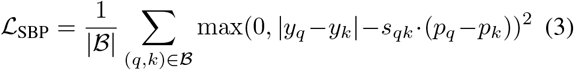

where *B* is a batch of node pairs sampled from the graph. By varying how we sample the node pair batches, we can make a model learn with focus on different aspects. We define a *local* and *global* version of the SBP loss corresponding intuitively to two commonly used error metrics in genome phasing, namely the Hamming and switch error.

### Global SBP loss

For each node *q* in the graph, we randomly select another node *k* to form a pair (*q, k*) for the contrastive loss computation. To do this efficiently in parallel, we convert the full set of nodes in the graph into a list and apply a random permutation, ensuring that each node appears exactly once as a query node, paired with a randomly selected key node.

### Local SBP loss

For each node *q*, we randomly sample a node *k* sharing an overlap edge with *q*. These represent sequences that are located nearby along the chromosome and incentivize locally coherent predictions.

The SBP loss exhibits distinct behavior depending on the label relationship. When both nodes have the same label (*y*_*q*_ = *y*_*k*_), the loss becomes (*p*_*q*_ −*p*_*k*_)^2^, directly penalizing any difference with no margin tolerance. When nodes have different labels (*y*_*q*_ ≠ *y*_*k*_), the loss encourages predictions to be separated by at least*| y_q_ y_k_ |*, with violations penalized quadratically. This formulation naturally guides the model toward the desired clustering structure while leveraging all label information available in our supervised problem setting. In analogy to the pair losses, we also create a supervised binary triplet (SBT) loss (detailed in Appendix A). We use an additional auxiliary loss to push predictions of nodes shared between clusters towards zero using a simple mean squared error. This can help the model converge earlier during training:

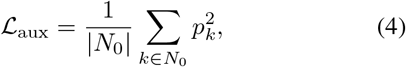

Where

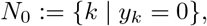

and *pk* are the corresponding predictions.

## 5. Model Architecture

Given an input graph with adjacency matrix **A** and node feature matrix **X***∈***ℝ***n*^×*d*^, the features are first embedded into an initial node embedding **Z**^(0)^ via a feed-forward inputlayer *f*in:

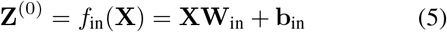

`grapHiC’s` model architecture is based on the simplified graph transformer (SG-Former) (Wu et al., 2024), consisting of a GNN module and a linear attention (LA) module. The GNN captures local graph topology, while the attention models global interactions. The final representation is:

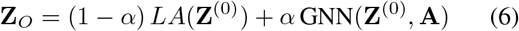

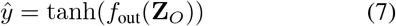

where *α* is a hyperparameter balancing the GNN and attention representations, *LA*(**Z**^(0)^) is the linear attention module, and *f*_out_ is a final feed-forward layer.

The GNN is implemented as a heterogeneous GIN (Xu et al., 2018) to handle multiple edge types *t∈ { o, h}* (overlap and Hi-C) with edge-separated, weighted adjacency matrices **A***t*. A single heteroGIN layer updates node embeddings as:

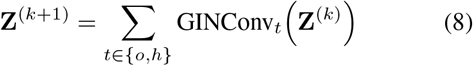

where each GINConv_*t*_ computes:

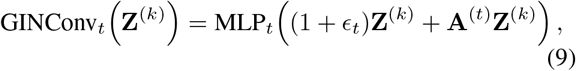

with learnable scalar *ϵ*_*t*_. The full GNN module GNN(**Z**^(0)^) denotes a stack of *l* such heteroGIN layers with residual connections and normalization.

This design allows the model to capture relation-specific patterns while properly weighting Hi-C edges by contact strength. Multiple heteroGIN layers are combined with global-attention-based embeddings in the prediction head, preserving both local graph topology and global interactions.

## 6. Hi-C enhanced Unitig Graphs

### 6.1. Data Pipeline

We construct Hi-C–enhanced unitig graphs from a set of simulated HiFi and real Hi-C reads from the same genome. HiFi reads are sampled from the haplotype-resolved I002C genome sequence using `PBSim3`(Sarashetti et al., 2024b). For the Hi-C reads, we use Omni-C reads used in the original genome assembly. `PBSim3` aims to emulate the error profile observed in PacBio Hi-Fi-based genome sequencing; using simulated data allows us to have ground truth position and haplotype information for each read, which we can use as labels for training and validation. To increase diversity across training graphs, we use different replicates of the Hi-C experiments and random subsamples of the simulated HiFi read for each graph. Unitig graphs were generated from HiFi reads using `hifiasm` v0.25 (Figure 2 **A**), then the Hi-C connections between unitigs are identified and normalized (Figure 2 **B**).

**Figure 2.**
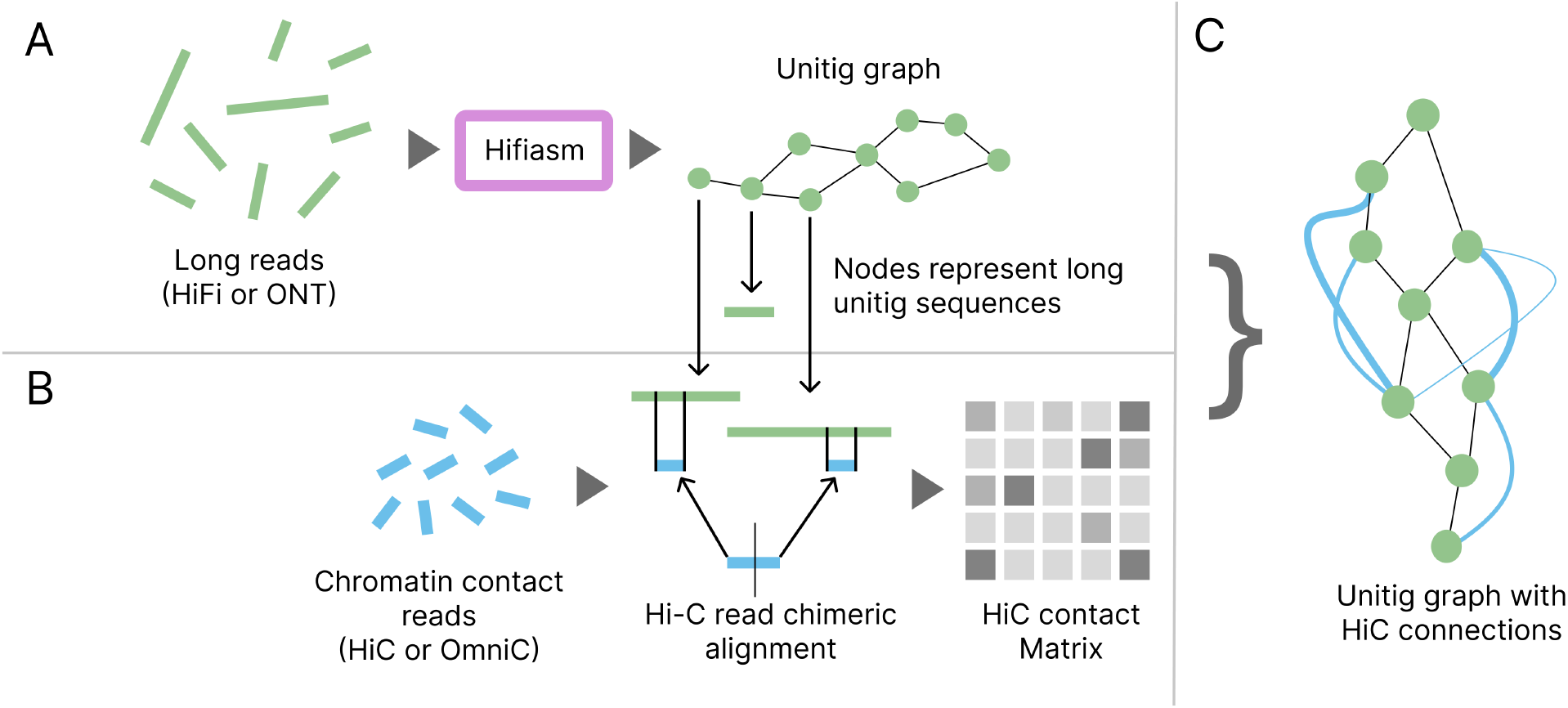
**A:** Long reads are processed by `hifiasm` to create a unitig graph. **B:** Chromatin Contact (Hi-C) reads are mapped against the unitigs, to create a contact matrix. **C:** The Contact matrix is realized as edges in the unitig graph, where the weight is defined by the amount of Hi-C contacts of unitigs.

**Figure 3.**
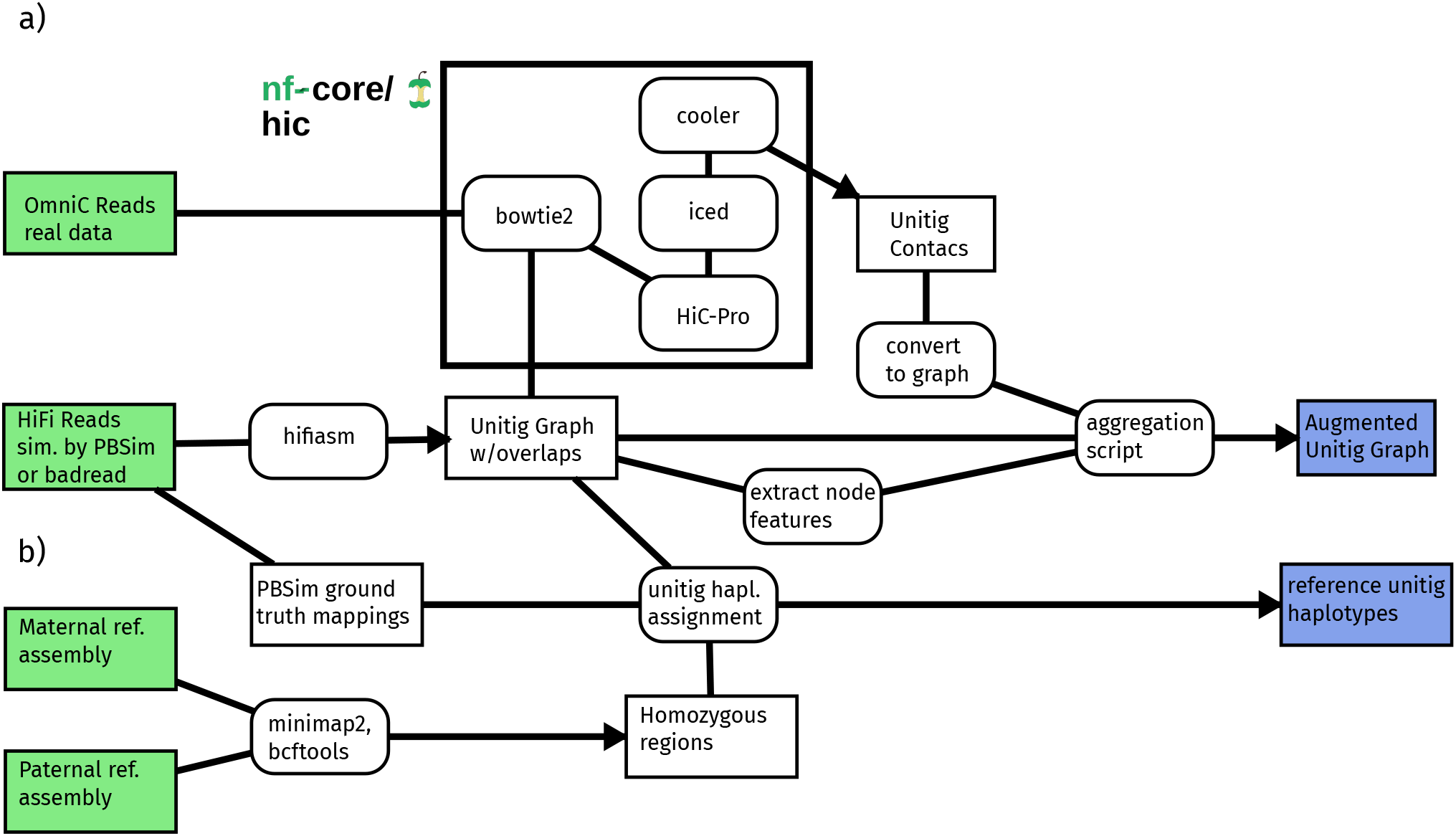
Pipeline used for dataset construction. Tools are shown in boxes with rounded edges, while intermediate outputs are shown in rectangular boxes. Initial inputs are highlighted in light green; outputs used for model training and evaluation are shown in light blue. **a)** To construct the augmented unitig graph, we generate unitigs with `hifiasm` on reads simulated with PBSim (PacBio HiFi) or `badread`(ONT). OmniC reads are mapped to the unitigs using the nf-core/hic pipeline based on the HiC-Pro toolchain. Contacts obtained from this process are added as additional edges to the existing overlap graph, with weights derived from ICE-normalized Hi-C contact frequencies. **b)** To obtain ground-truth haplotype information for phasing loss calculation, we combine the read simulator’s origin loci for all reads within each unitig. Additionally, homozygous regions are identified by variant calling between the assembled parental genomes and subsequently annotated.

**Figure 4.**
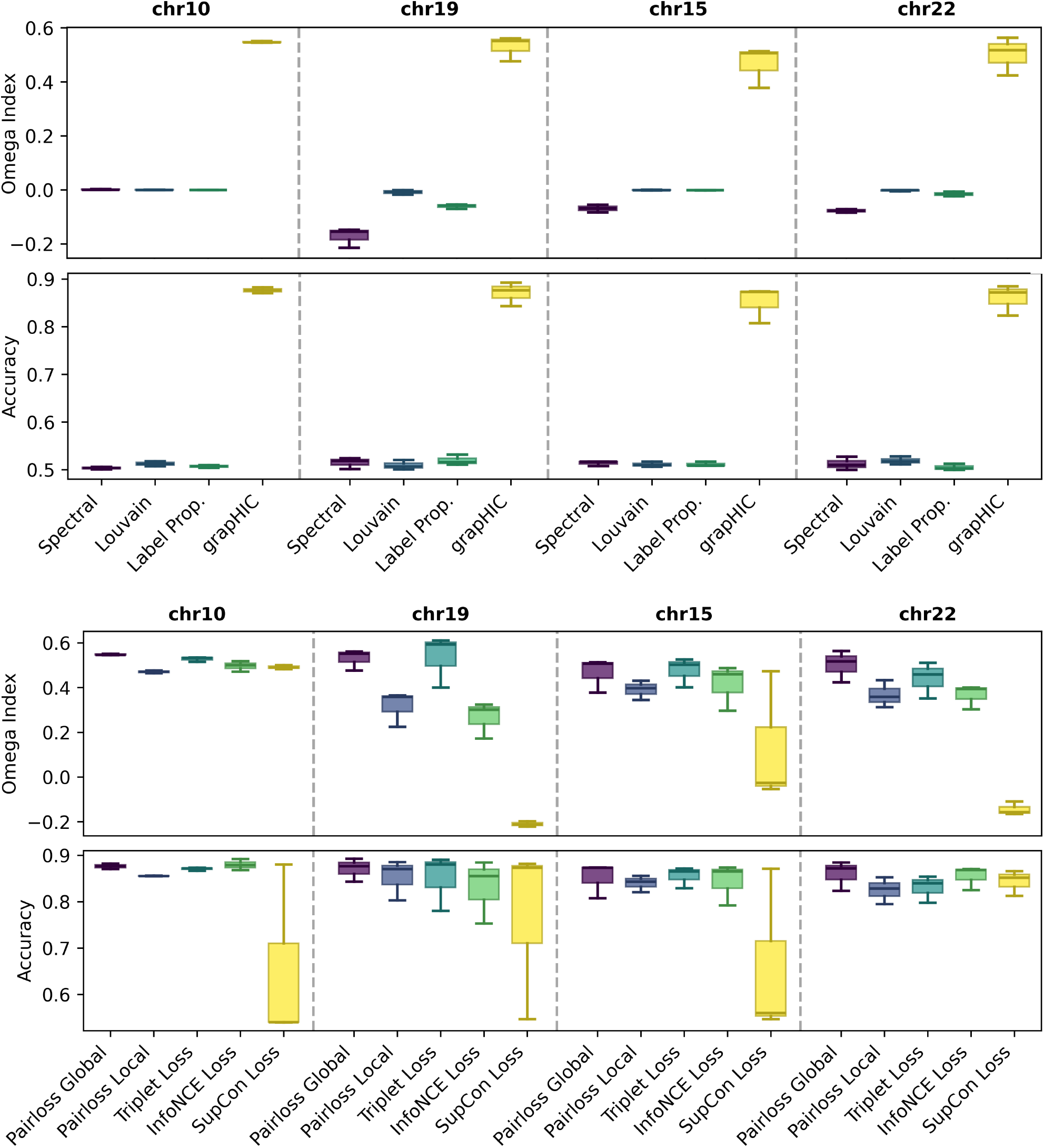
Comparing `grapHiC` with 1) classic clustering algorithms and 2) comparison of differnet contrastive learning functions. Each chromosome is evaluated with three different independently sampled graphs. Note that Chrs. 15 and 22 are acrocentric. The Tukey plot shows the minimum, maximum and median sample as well as standard deviation of the three samples. Accuracy is computed from the best one-to-one mapping as determined by the Hungarian Method between predicted cluster labels and true labels. Omega Index is an extension of Adjusted Random Index (ARI) for fuzzy clustering. The Omega index is an extension of the ARI to fuzzy clustering, and can take values from −1 to +1, with a perfect clustering being assigned +1 and a random assignment 0. An accuracy of 0.5 corresponds to random cluster assignments, while 1.0 would mean perfect clustering.

**Figure 5.**
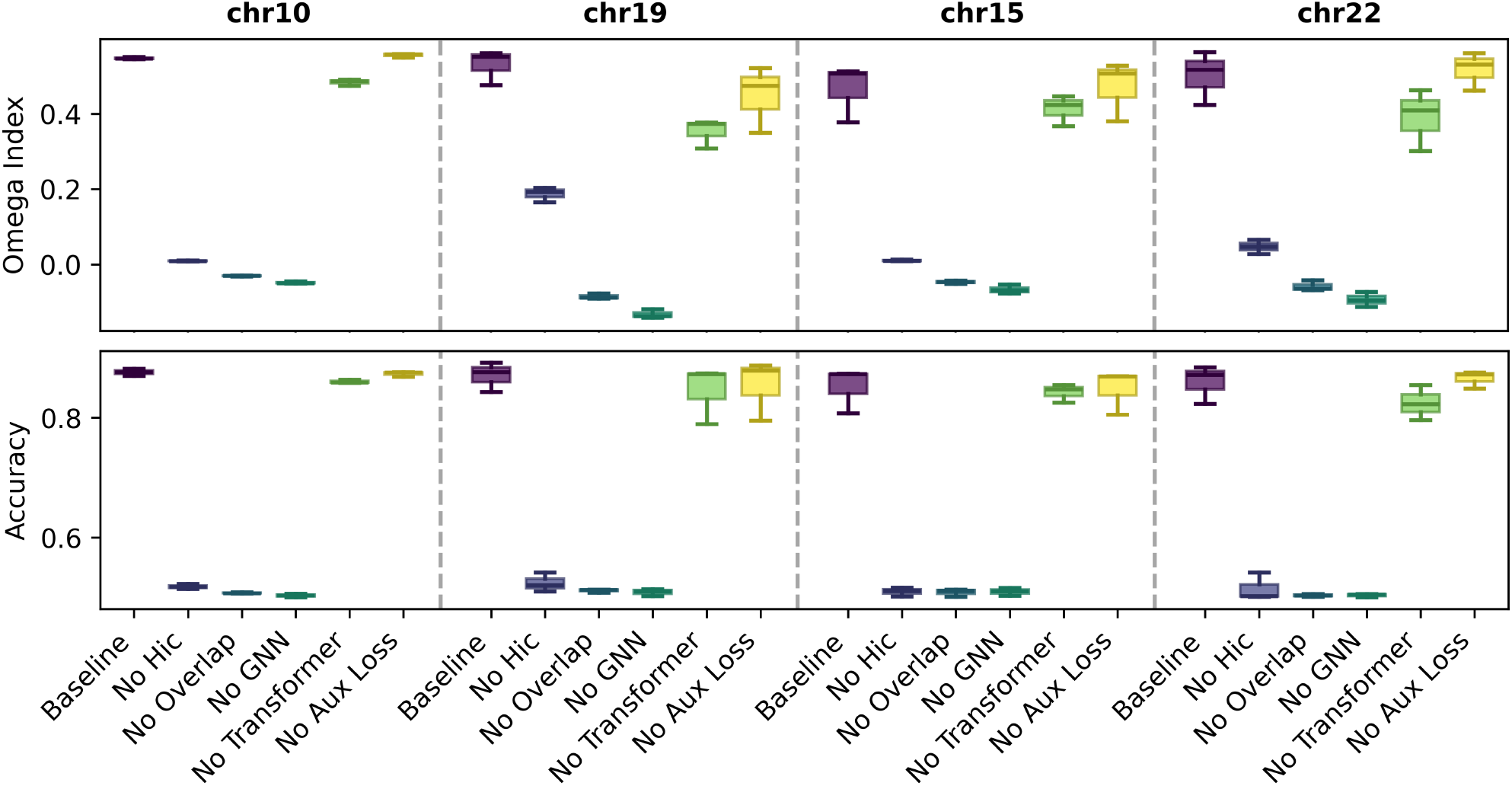
Ablation study for grapHIC. Each chromosome is evaluated with three different graphs sampled from different sets of HiFi reads. The Tukey plot shows the minimum, maximum and median sample as well as the standard deviation of the three samples.

For this mapping step, we implement two alternative pipelines: an **accurate** pipeline and a **fast** pipeline. The accurate pipeline is based on the nf-core/hic pipeline and uses base-level alignment with bowtie2 (Langmead & Salzberg, 2012) followed by normalization using `Cooler` (Abdennur & Mirny, 2019). The fast pipeline replaces exact alignment with a minimizer-based mapping using `minimap2` (Cheng et al., 2021) and applies symmetric adjacency normalization. See Appendix B for detailed descriptions of both pipelines. The output of the data preparation pipeline is an augmented unitig graph containing both sequence overlaps and Hi-C contacts between unitigs as edges. Hi-C edges have a weight indicating the strength of the connection between the nodes (Figure 2 **C**). Nodes in the graph have several features: the number of adjacent overlaps and Hi-C edges, the length of the sequence of the unitig, and the amount of reads supporting the unitig as reported by `hifiasm`. Details of the feature extraction are given in Appendix C.

### 6.2. Dataset

We train our model on two synthetic datasets prepared from synthetic reads sampled to 40x coverage from the I002C phased human genome (Sarashetti et al., 2024b): **Dataset (I)** is a single chromosome dataset created with the **accurate** pipeline consisting of 48 graphs containing of one chromosome each. **Dataset (II)** is a dataset created with the **fast** pipeline consisting of 35 full-genome graphs containing all autosomal chromosomes. See Appendix D for more information on the dataset, Appendix E for details of `graphic`’s training and Appendix F for used computational resources.

## 7. Evaluation: Clustering

To evaluate the clustering capability, we compare accuracy (of the best one-to-one mapping between predicted cluster labels and true labels, using the Hungarian method), Adjusted Rand Index (ARI) and Normalized Mutual Information (NMI)(Hubert & Arabie, 1985; McDaid et al., 2013). Since these methods can only evaluate disjoint clustering, we also include the Omega Index (Collins & Dent, 1988), an extension of the ARI for non-disjoint (fuzzy) solutions. For evaluation, we show `graphic`’s performance on the four datasets compared to three traditional clustering algorithms. We compare `graphic` to spectral clustering (Ng et al., 2001), the Louvain algorithm (Blondel et al., 2008), and Label Propagation (Zhu & Ghahramani, 2002) as alternative approaches to cluster nodes into haplotypes.

We compare different versions of `graphic` trained on **Dataset (I)** using different objective functions and evaluate their performance against our proposed SBP-Loss. We include both SBP-Loss (global) and SBP-Loss (local), as well as the SBT-Loss all described in Section 4. We compare our losses to a supervised version of InfoNCE (Oord et al., 2018) and the Supervised Contrastive Loss (SupCon) (Khosla et al., 2020), each with a 64-dim embedding output and an additional 16-dim projection head that is not used during evaluation. Similarly to our proposed losses, both of these approaches use batchwise processing of 16 negative (and in the case of SupCon also 16 positive) samples. The clustering is performed using *k*-means clustering for Accuracy, NMI, and ARI metric computation and Fuzzy *c*-Means clustering (Bezdek et al., 1984) for the Omega Index computation. Table 1 shows the Accuracy and Omega Index of the compared methods. Appendix G shows the results graphically and adds NMI and ARI results.

**Table 1.** Comparison of different phasing methods on the chromosome validation sets. Accuracy and Omega Index reported are averaged for all replicates per chromosome. Best scores are marked in bold.

| Method | chr10 |  | chr19 |  | chr15 |  | chr22 |  |
| --- | --- | --- | --- | --- | --- | --- | --- | --- |
|  | Acc | Omega | Acc | Omega | Acc | Omega | Acc | Omega |
| Spectral | 0.503 | 0.002 | 0.515 | -0.172 | 0.514 | -0.069 | 0.513 | -0.077 |
| Louvain | 0.513 | 0.000 | 0.510 | -0.009 | 0.511 | -0.001 | 0.519 | -0.002 |
| LP | 0.507 | -0.000 | 0.520 | -0.061 | 0.511 | -0.001 | 0.505 | -0.015 |
| SBP Loss (global) | 0.876 | **0.548** | **0.871** | **0.530** | 0.852 | **0.466** | **0.860** | **0.501** |
| SBT Loss | 0.871 | 0.527 | 0.850 | 0.535 | **0.855** | 0.476 | 0.831 | 0.441 |
| SBP Loss (Local) | 0.855 | 0.470 | 0.853 | 0.316 | 0.840 | 0.391 | 0.825 | 0.368 |
| SupCon | 0.653 | 0.491 | 0.767 | 0.210 | 0.659 | 0.131 | 0.843 | 0.143 |
| InfoNCE | **0.880** | 0.496 | 0.831 | 0.266 | 0.844 | 0.414 | 0.855 | 0.366 |

We conduct an ablation study to assess the impact of each component of `graphic` on prediction quality. Overlap and HiC-edges, as well as the GNN component, are all required for better-than-random performance by `grapHiC.` Removing the attention component or the auxiliary loss mildly decreases performance. Details are given in Appendix H.

## 8. Evaluation: Genome Assembly

To evaluate our model in a real-world use case, we test whether `graphic` predictions can be useful for *de novo* genome assembly. For this, we use the recently published assembler `DipGNNome` (Schmitz et al., 2026) and a `graphic` model trained on **Dataset (II)**. We use `grapHiC`to assign a haplotype score to each node in the graph. These scores are then used in the beam search process of `DipGNNome` to separate the haplotypes during the assembly process. Appendix I provides more details on how `grapHiC’s`predictions are used to find assemblies using `DipGNNome.`

We evaluate the combined `D-grapHiC (DipGNNome + grapHiC)` pipeline on synthetic full-genome samples generated by `PBsim3`(Ono et al., 2022), using read error profiles and length distributions matched to real HG002 HiFi reads. Additionally we use real experimental Hi-C reads, randomly downsampled to 50 million pairs for performance reasons. To assess the contribution of `graphic`, we also compare against `DipGNNome`alone.

The resulting assemblies are compared against those produced by `hifiasm` (Hi-C), which is particularly informative for two reasons. First, `hifiasm` represents the strongest state-of-the-art baseline for de novo assembly from single long-read technologies. Second, by using `hifiasm’s`overlap-based phasing to construct the unitig graphs, we can directly compare the layout and phasing algorithms—the steps that simplify the graph, resolve haplotypes, and assemble sequences—starting from the same input graph.

### Genome Assembly Experiments

We evaluate`;D-grapHiC`on two full-genome synthetic graph settings: (i) I002C, the genome used for training (but graphs created out of reads not seen during training), and (ii) HG002, a different high-quality human genome. Graphs generated from different read sets of the same genome can exhibit substantially different topologies and attributes. We therefore assess how performance varies across diverse graph topologies both within a single genome and across different human genomes.

Table 2 compares `DipGNNome, grapHiC + DipGNNome (D-grapHiC),` and `hifiasm` across multiple I002C and HG002 replicates. The results show that while DipGNNome achieves high contiguity, it fails to phase haplotypes in this setting, with Hamming error rates consistently around 40%. This behavior is expected, as `DipGNNome`relies on trio-binning markers for phasing, which are not available in our experimental setup.

**Table 2.** Comparison of DipGNNome, D-grapHiC, and Hifiasm on I002C and HG002 replicates. Bolded are the best metrics of only the comparsion **D-grapHiC** and **Hifiasm**.

| Dataset | Metric | DipGNNome | D-graphHiC | Hifiasm |
| --- | --- | --- | --- | --- |
| **I002C(1)** | Length (Mb) | 2861.3 / 2859.7 | 2692.4 / 2662.8 | 2779.1 / 2780.3 |
|  | Rdup (%) | 1.4 / 1.4 | **0.2 / 0.2** | 0.5 / 0.5 |
|  | NG50 (Mb) | 129.8 / 129.8 | **98.9 / 99.6** | 82.2 / 82.2 |
|  | Switch Err (%) | 0.7 / 0.7 | **0.3 / 0.5** | 0.4 / **0.5** |
|  | Hamming Err (%) | 40.2 / 40.0 | 1.2 / 1.6 | **0.7 / 0.8** |
| **I002C(2)** | Length (Mb) | 2865.4 / 2868.1 | 2685.3 / 2659.5 | 2785.3 / 2769.1 |
|  | Rdup (%) | 1.5 / 1.4 | **0.3 / 0.1** | 0.5 / 0.5 |
|  | NG50 (Mb) | 133.9 / 134.3 | **91.6 / 87.0** | 86.1 / 85.9 |
|  | Switch Err (%) | 0.7 / 0.7 | **0.3 / 0.5** | 0.4 / **0.5** |
|  | Hamming Err (%) | 40.2 / 39.5 | 1.3 / 1.8 | **0.9 / 1.0** |
| **I002C(3)** | Length (Mb) | 2877.9 / 2872.9 | 2688.0 / 2661.4 | 2764.8 / 2800.3 |
|  | Rdup (%) | 1.6 / 1.5 | **0.2 / 0.1** | 0.4 / 0.7 |
|  | NG50 (Mb) | 111.2 / 111.2 | **95.8 / 93.6** | 86.1 / 92.1 |
|  | Switch Err (%) | 0.6 / 0.6 | **0.3 / 0.5** | 0.4 / **0.5** |
|  | Hamming Err (%) | 39.1 / 39.3 | 1.1 / 2.1 | **1.0 / 0.9** |
| **HG002(1)** | Length (Mb) | 2968.4 / 2971.9 | 2746.3 / 2733.7 | 2881.2 / 2854.0 |
|  | Rdup (%) | 1.6 / 1.3 | **0.2 / 0.2** | 0.7 / 0.5 |
|  | NG50 (Mb) | 137.5 / 134.7 | **98.7 / 86.9** | 90.0 / **89.6** |
|  | Switch Err (%) | 1.5 / 1.5 | 1.8 / **1.1** | **1.4 / 1.5** |
|  | Hamming Err (%) | 40.7 / 40.8 | 3.2 / 2.7 | **2.1 / 1.9** |
| **HG002(2)** | Length (Mb) | 2966.8 / 2971.4 | 2724.6 / 2736.0 | 2868.8 / 2866.1 |
|  | Rdup (%) | 1.7 / 1.6 | **0.2 / 0.1** | 0.6 / 0.8 |
|  | NG50 (Mb) | 119.8 / 121.2 | **91.2 / 72.3** | 89.0 / **86.5** |
|  | Switch Err (%) | 1.5 / 1.5 | 1.7 / **1.1** | **1.5 / 1.3** |
|  | Hamming Err (%) | 39.3 / 40.8 | 3.9 / 3.1 | **1.9 / 1.6** |
| **HG002(3)** | Length (Mb) | 2959.4 / 2966.0 | 2748.1 / 2729.8 | 2872.5 / 2866.5 |
|  | Rdup (%) | 1.5 / 1.5 | **0.2 / 0.1** | 0.7 / 0.6 |
|  | NG50 (Mb) | 134.1 / 133.7 | 87.3 / 76.2 | **89.8 / 86.7** |
|  | Switch Err (%) | 1.5 / 1.5 | 1.7 / **1.1** | **1.4 / 1.4** |
|  | Hamming Err (%) | 41.4 / 41.6 | 3.1 / 3.6 | **3.0 / 2.5** |

In contrast,`D-grapHiC,` which uses `grapHiC`predictions to guide phasing within `DipGNNome`;, demonstrates that the learned phasing scores can effectively resolve haplotypes. Across all datasets, the Hamming error is reduced substantially, to values between 1.1% and 3.9%. This confirms that `graphic`;provides a reliable phasing signal that can be directly integrated into an OLC-style assembler.

On both the synthetic I002C and HG002 datasets, `D-grapHiC`achieves overall performance comparable to `hifiasm,` the current state-of-the-art diploid assembler for HiFi long-read data. Phasing accuracy measured by switch error is very similar between the two methods: `hifiasm`shows slightly lower switch error for three haplotypes, while `D-grapHiC` performs slightly better for nine haplotypes. Hamming error rates for D-grapHiC are generally higher. Notably, higher Hamming error rates relative to `hifiasm`were also reported in the original DipGNNome study (Schmitz et al., 2026), suggesting that this difference is not primarily attributable to grapHiC’s phasing predictions.

In terms of contiguity, `D-grapHiC` assemblies achieve NG50 values comparable to those of hifiasm. Duplication rates are consistently lower for `D-grapHiC` across all datasets. Overall, these results demonstrate that grapHiC-guided assemblies produced by `DipGNNome` can match state-of-the-art performance in both contiguity and phasing accuracy, while in some cases reducing the rate of erroneous duplications. The reduced switch error and fewer erroneous duplications may reflect D-grapHiC’s ability to avoid contig misjoins by phasing prior to graph simplification.

## 9. Conclusion

We present `graphic`a machine-learning framework for reference-free haplotype phasing on unsimplified unitig graphs. It integrates Hi-C and overlap information, distinguishes homozygous from heterozygous regions, and is trained using the Supervised Binary Pair Loss, which accounts for label-switching symmetry and improves both accuracy and training stability. On synthetic and human datasets,`graphic` combined with `DipGNNome`produces well-phased assemblies with low switch and Hamming error rates and contiguity comparable to the state of the art. Importantly`graphic` can be retrained on any graph dataset, enabling integration with different sequencing technologies and assembly approaches.

This proof-of-concept evaluation is restricted to synthetic long reads combined with experimental Hi-C data on human genomes. Extending the benchmarking to full real-world, non-human datasets is an important direction for future work.

An especially promising direction is the extension to polyploid genomes, where disentangling multiple haplotypes poses an even greater challenge largely unaddressed by most classical and all Machine Learning-based phasing approaches. In these cases, the strength of grapHiC of being capable of phasing prior to simplification and its potential to avoid misjoining unitigs from different haplotypes could improve significantly upon the state of the art.

## A. Supervised Binary Triplet Loss

### SBT Loss

We create a supervised binary triplet (SBT) loss, based on the triplet loss common in contrastive learning. For this, we consider triplets of samples (*q, kp, k_n_*) where *q* is the query/anchor sample, *k*_*p*_ is a positive key sample (same class as query), and *k*_*n*_ is a negative key sample (different class from query). The general triplet loss can be formulated as:

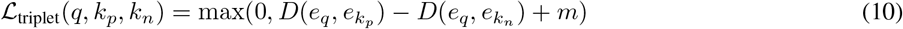

Where *D* is a distance function, usually the squared Eucledian distance. *eq, ek_p_*, and *ek*_*n*_ are the embeddings of the query, positive key, and negative key samples respectively, and *m* is a margin parameter. In our binary haplotype phasing context, using one-dimensional embeddings, and batches over all nodes in the graph, we formulate the loss as:

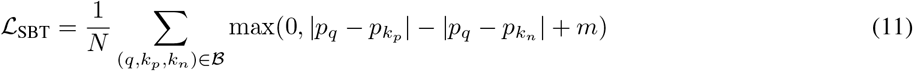

Where *pq, pk_p_*, and *pk*_*n*_ are the scalar predictions for the query, positive key, and negative key samples. For finding query samples, we select random nodes from the set of desired classes, where nodes of shared clusters are added to the positive set. (Note that this takes longer than the selection of samples in pair loss (global), where we only need to shuffle the list of nodes once and use it for the whole batch). The margin *m* is set to 1.

## B. Details: Hi-C Alignment Pipelines

### B.1. Accurate Hi-C Alignment Pipeline

To accurately align Hi-C reads to the unitig graph, we use the nf-core/hic pipeline (Servant et al., 2023), which is based on HiC-Pro (Servant et al., 2015). A high-level overview of the dataset preparation procedure is shown in Suppl. Figure 3.

The pipeline first performs read quality control with FastQC and MultiQC (Ewels et al., 2016), then aligns the reads to the unitigs using bowtie2 (Langmead & Salzberg, 2012). Unlike the standard practice in Hi-C analysis for chromosomal contact discovery, we use sensitive local alignment, since the unitigs may still contain sequencing errors that are only corrected in the final assembly stage. A custom script employing Cooler (Abdennur & Mirny, 2019) aggregates Hi-C connections per unitig.

### B.2. Approximate Hi-C Mapping Pipeline

The approximate mapping pipeline replaces base-level alignment with a minimizer-based approach. Specifically, Hi-C reads are mapped to unitigs using minimap2 instead of bowtie2, and ICE normalization via Cooler is replaced with symmetric degree normalization of the adjacency matrix.

The adjacency matrix *A* with Hi-C contact info asedge weights is symmetrically degree-normalized:

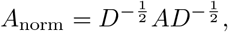

where *D* is the diagonal degree matrix with entries *Dii* = _*j*_ *Aij*. Below is the detailed command used to map Hi-C reads against unitigs:

~~~
# Mapping Hi-C reads against unitigs using minimap2
 minimap2 -x sr -N 10 -p 0.8 -t <threads> \
 <unitigs.fasta> <hic_R1.fastq.gz> <hic_R2.fastq.gz> \
 > alignments.paf
~~~

Here, `-x sr`configures `minimap2` for short-read mapping (appropriate for Hi-C data), `-N 10`outputs up to ten secondary alignments per read, and `-p 0.8` specifies the minimum identity threshold. We then parse the PAF file and generate a weighted contact edge list with a custom script. Contact edges are then weighted by the number of supporting read pairs.

#### B.3. Data Pipeline Output Graphs

The output of the data preparation pipeline is a heterograph where nodes represent unitigs and edges represent either sequence overlaps or Hi-C contacts. Each node is annotated with features extracted during preprocessing, and edges are labeled by type and assigned weights.

The full graph is serialized into the .`pyg` format using `networkx`and custom scripts. This makes it directly consumable by PyTorch Geometric-based models.

## C. Node and Edge features

### Node features

Each unitig node has four primary features:

- **Sequence length** Computed from the FASTA sequences of the unitigs and divided by 10,000 to scale values into a range typically within [0, 10].
- **Coverage** Read coverage is extracted directly from the unitig metadata in the GFA output of hifiasm. Specifically, each S-line includes the number of reads supporting the unitig, which we use as an unnormalized scalar feature.
- **Degree** The degree or number of adjacent Hi-C and overlap edges are each recorded as separate features, capturing how connected each unitig is in the graph under each edge type.

### Edge features

Edges are annotated with both a type label (`either overlap` or `hic`) and a weight:

- **Overlap edges:** These represent exact or near-exact overlaps between unitigs as derived from hifiasm. All overlap edges are assigned a constant weight of 1.
- **Hi-C edges:** These reflect long-range physical proximity contacts inferred from Hi-C sequencing. The weight is the normalized contact entry between the two connected unitigs.

## D. Dataset Details

### D.1. Dataset (I): Single Chromosome

We evaluate our model on three synthetic datasets consisting of synthetic reads sampled to 40x coverage from the I002C phased human genome (Sarashetti et al., 2024b). We use PBSim3 (Ono et al., 2022) with flag *–method sample* to mimic read length and error distribution of the real experimental I002C reads.

Each dataset includes all autosomes except Chromosome 14. We choose two non-acrocentric chromosomes as validation datasets (2 *×* 3 graphs) and two acrocentric as well as two non-acrocentric chromosomes as test datasets (4 *×* 3 graphs), and the rest for training (16 *×* 3 graphs, containing the other two acrocentrics). Acrocentric chromosomes contain the long and complex rDNA repeats encoding the ribosomal RNAs that are challenging to assemble and phase, and thus represent a particularly tough challenge for our model. The four graphs in the test set are Chromosome 10 (medium size, non-acrocentric), Chromosome 15 (medium size, acrocentric), Chromosome 19 (short, non-acrocentric) and Chromosome 22 (short, acrocentric).

The training dataset consists of 45 graphs with a total of 139,574 nodes (mean: 3,101.64, std: 1,144.56) ranging from 982 to 5,427 nodes in each graph, where each unitig corresponds to tens to hundreds of thousands of base pairs of DNA. The graphs contain in total 1,043,808 (mean: 23,195.73, std: 31,544.18) sequence overlap edges, with a range of 1,355 to 107,087 in each graph and in total 2,584,249 (mean: 57,427.76, std: 28,061.06) HiC-connections with a range of 9,314 to 101,487 in each graph.

#### D.1.1. Dataset (II): Full Genome

For training the new `grapHiC` model, we use and augment the dataset presented in `DipGNNome` (Schmitz et al., 2026), consisting of 35 graphs based on the I002C (human) genome. To introduce variation, we randomly subsample 50 million Hi-C read pairs from the original dataset for each graph. The resulting graphs combine real Hi-C data with synthetic HiFi data generated using `PBSIM3`. We split the dataset into 30 graphs for training and 5 for validation.

## E. Training Setup

We train grapHIC using a training pipeline implemented in PyTorch, with PyTorch-Geometric.

Table 3 summarizes the key training parameters.

**Table 3.** SGFormer Training Parameters.

| Parameter | Value |
| --- | --- |
| GCN Layers | 8 |
| Transformer Layers | 3 |
| Hidden Dimension | 256 |
| Learning Rate | $10^{-4}$ |
| Auxiliary Loss Weight | 0.02 |
| Layer Normalization | True |
| GNN Dropout | 0.1 |

**Table 4.** Comparison of clustering methods by chromosome using NMI and ARI metrics. Best scores in each column are highlighted in bold. SCP Loss (Global) and InfoNCE demonstrate superior performance across chromosomes.

| Method | chr10 |  | chr19 |  | chr15 |  | chr22 |  |
| --- | --- | --- | --- | --- | --- | --- | --- | --- |
|  | NMI | ARI | NMI | ARI | NMI | ARI | NMI | ARI |
| Spectral | 0.000 | 0.000 | 0.003 | 0.000 | 0.003 | 0.000 | 0.002 | 0.000 |
| Louvain | 0.001 | 0.000 | 0.001 | 0.000 | 0.000 | 0.000 | 0.001 | 0.000 |
| LP | 0.000 | 0.000 | 0.002 | 0.001 | 0.000 | 0.000 | 0.000 | -0.001 |
| SCP Loss (global) | 0.469 | 0.567 | **0.451** | **0.551** | **0.407** | 0.498 | **0.438** | **0.521** |
| SBT Loss | 0.448 | 0.549 | 0.411 | 0.501 | 0.409 | **0.506** | 0.378 | 0.439 |
| SCP Loss (Local) | 0.414 | 0.505 | 0.410 | 0.503 | 0.377 | 0.463 | 0.369 | 0.425 |
| SupCon | 0.185 | 0.197 | 0.321 | 0.383 | 0.172 | 0.191 | 0.433 | 0.473 |
| InfoNCE | **0.479** | **0.577** | 0.383 | 0.451 | 0.396 | 0.478 | 0.406 | 0.504 |

**Table 5.**
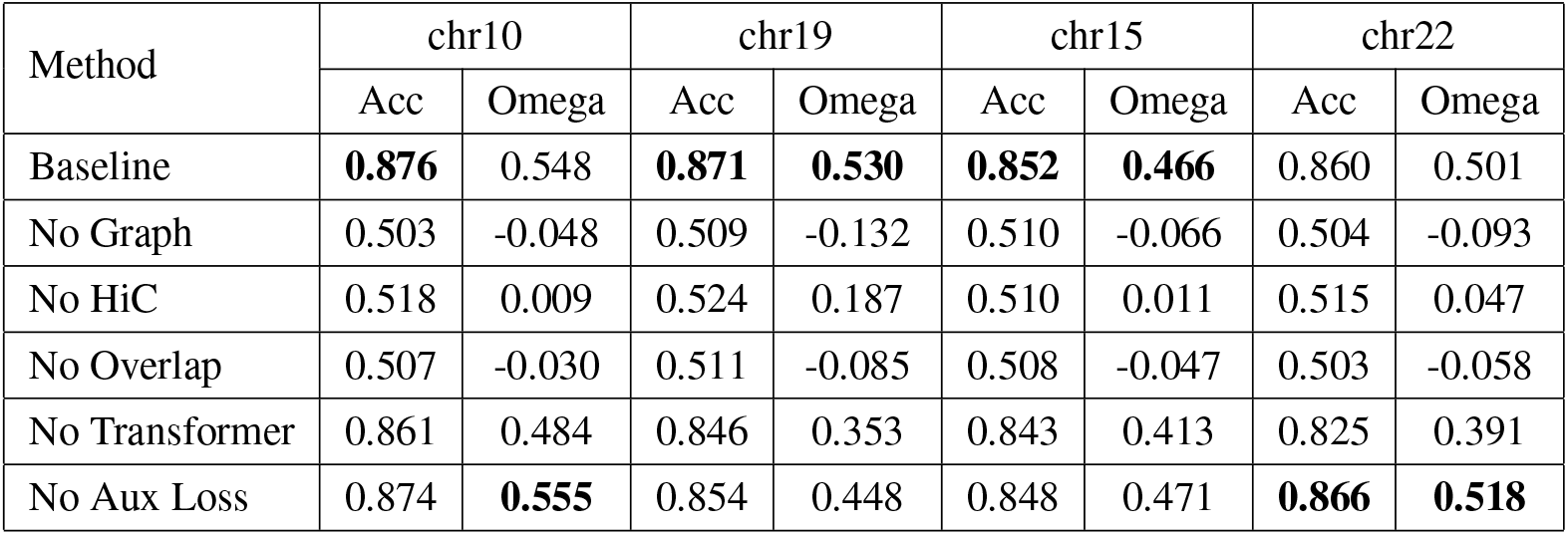
Performance comparison of different methods across chromosomes. The table shows accuracy (Acc) and Omega Index values for each method on chromosomes 10, 19 (non-acrocentric), 15, and 22 (acrocentric). Bold values indicate the best performance for each chromosome.

| Method | chr10 |  | chr19 |  | chr15 |  | chr22 |  |
| --- | --- | --- | --- | --- | --- | --- | --- | --- |
|  | Acc | Omega | Acc | Omega | Acc | Omega | Acc | Omega |
| Baseline | **0.876** | 0.548 | **0.871** | **0.530** | **0.852** | **0.466** | 0.860 | 0.501 |
| No Graph | 0.503 | -0.048 | 0.509 | -0.132 | 0.510 | -0.066 | 0.504 | -0.093 |
| No HiC | 0.518 | 0.009 | 0.524 | 0.187 | 0.510 | 0.011 | 0.515 | 0.047 |
| No Overlap | 0.507 | -0.030 | 0.511 | -0.085 | 0.508 | -0.047 | 0.503 | -0.058 |
| No Transformer | 0.861 | 0.484 | 0.846 | 0.353 | 0.843 | 0.413 | 0.825 | 0.391 |
| No Aux Loss | 0.874 | **0.555** | 0.854 | 0.448 | 0.848 | 0.471 | **0.866** | **0.518** |

**Table 6.** Beam search algorithm configuration and heuristic parameters.

| General Parameters |  |
| --- | --- |
| Beam width (k) | 5 |
| Samples (n) | 100 |
| Min. contig length ($P_{\min}$) | 100k |
| Min. component length ($C_{\min}$) | 25 |
| Beam Heuristic Parameters |  |
| **grapHiC + DipGNNome** |  |
| $\alpha$ | $10^{-4}$ |
| $\beta$ | 10 |
| $\gamma$ | 20 |
| $c_b$ | $> 0.9$ |
| $c_t$ | $> 0.5$ |

For optimization, we use the Adam optimizer with a learning rate of 10^−4^. The training process supports multiple loss functions, including global pairwise loss, local pairwise loss, and triplet loss for prediction-based approaches, as well as contrastive embedding losses (SupCon and InfoNCE) for embedding-based approaches. An auxiliary loss with a weight of 0.02 is applied to encourage shared embeddings to approach zero.

During training, we evaluate model performance using the validation loss. We let the models run for 2000 epochs and select the one with the lowest loss on the validation dataset (see 6.2). The training configuration includes dropout regularization and layer normalization in each GNN layer block. We use Weights & Biases (Biewald, 2020) for experiment tracking, logging metrics such as loss values and accuracies.

## F. Compute Resources

All runs of the dataset preparation pipeline were performed on two nodes with 2 TB of RAM and 64 cores in 3-5 days. The pipeline was configured so each run would use at most 512 GB of RAM and 16 cores by setting Nextflow’s job limits to the appropriate value. These steps, along with the conversion of the hifiasm output to a graph representation, also account for the major performance bottlenecks in the pipeline. The latter step involves a lot of memory operations in a Python script and presents an opportunity for runtime improvements even after considerable optimization of the implementation. In particular, meaningful parallelization could speed this task up immensely.

Because the pipeline is implemented in Nextflow, it natively supports most common compute infrastructures like local execution, AWS, Google Cloud, and a variety of HPC scheduling systems. The nf-core parts of the pipeline support all container architectures used by the nf-core project (docker, singularity, and conda environments). The aggregation steps at the end of the pipeline require a conda environment containing Cooler (Abdennur & Mirny, 2019) and NetworkX(Hagberg et al., 2008).

Execution of the pipeline requires considerable scratch space (up to a few hundred GB), as temporary read alignment files are written to disk multiple times if the Hi-C read alignment is broken up into chunks to reduce memory usage.

## G. Clustering Results Details

This section shows the clustering experiment results in more detail. Suppl. Figure 4 shows the comparison of `grapHiC` against classical clustering methods graphically. Table **??**additionally shows the ARI and NMI metrics for the comparison of different contrastive learning losses used to train `grapHiC`.

## H. Ablation Study

Suppl. Figure 5 shows our ablation study. We can show that each component is necessary to get best results with `grapHiC.`

## I. DipGNNome + grapHiC details

Figure 6 illustrates the combined approach of `DipGNNome` and `graphic` called `D-grapHiC` which we use to obtain the results in our experiments.

**Figure 6.**
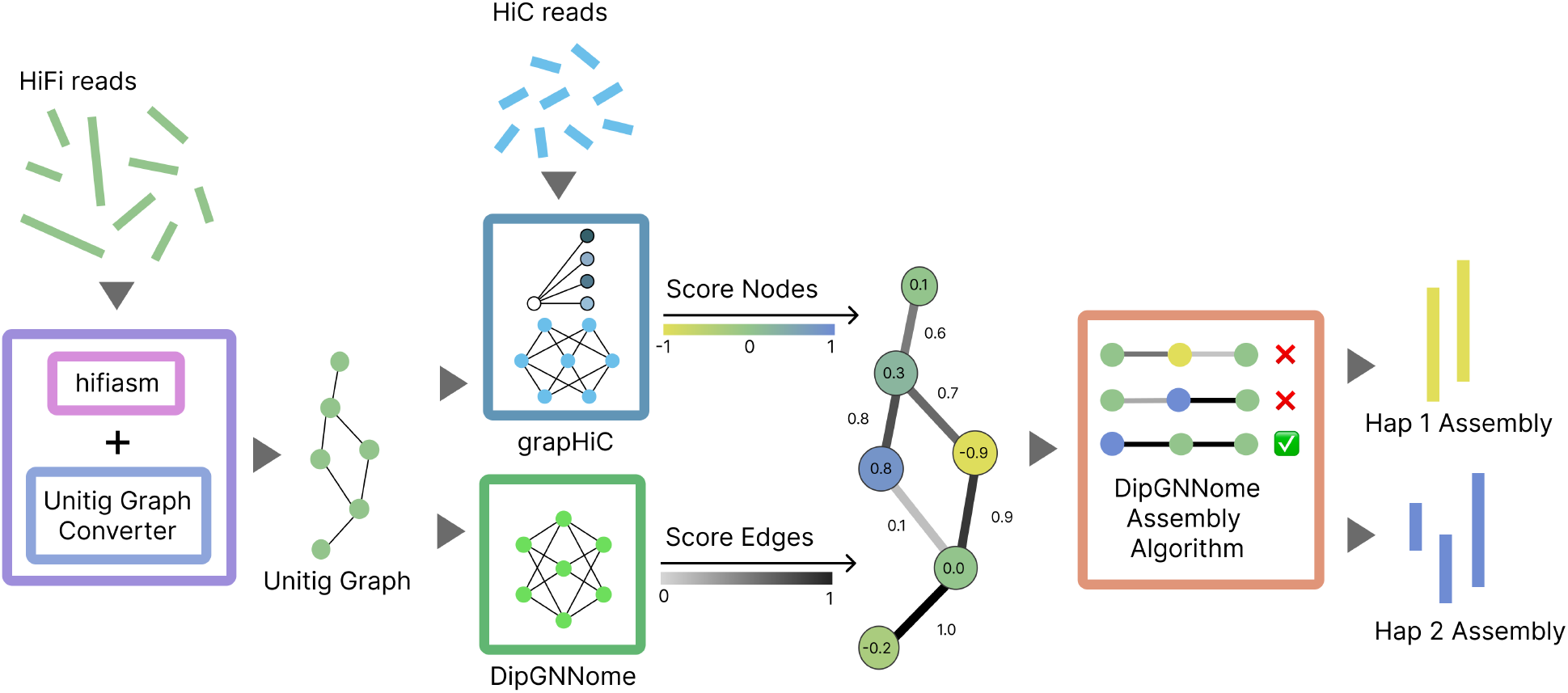
The `D-grapHiC` pipeline begins by constructing a unitig graph from HiFi reads using `hifiasm` together with the `DipGNNome` unitig processor. `DipGNNome`’s GNN then scores the edges of the unitig graph. In parallel, grapHiC maps Hi-C reads against the unitigs and assigns node scores representing haplotype probabilities. Finally, a `DipGNNome`-based assembly algorithm integrates the edge scores from DipGNNome and the node scores from `graphic` to generate a dual-haplotype assembly.

The original beam-guiding heuristic of `DipGNNome:`

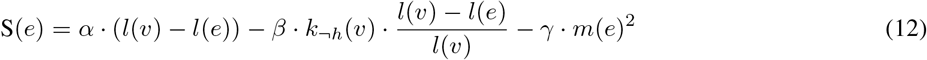

where *l*(*v*) is the length of node *v, l*(*e*) is the length of edge *e, k_h_*(*v*) is the number of k-mers not matching the target haplotype, and *m*(*e*) is the model prediction score for edge *e*.

To replace parental information with `grapHiC’s` HiC-based predictions, we simply replace the non-matching k-mer counts with a penalty based on `grapHiC` predicted haplotype.

Without loss of generality, we describe assembling haplotype 1. For this case, we interpret the continuous prediction score *p*(*v*) *∈* [***−***1, +1] from `graphic` as:

- *p*(*v*) *> ϵ*: predicted haplotype 1
- *p*(*v*) *<* ***−****ϵ*: predicted haplotype 2
- *|p*(*v*)*| ≤ ϵ*: homozygous

We determine the optimal value of *ϵ* using the validation set of `grapHiC`. Since the output layer of `grapHiC` uses a tanh activation, the predictions *p* lie in the range [***−***1, 1]. For each trained model, we evaluate *ϵ* values in the interval [0, 1] with a step size of 0.01 and select the value that achieves maximal accuracy in distinguishing homozygous from heterozygous nodes.

We then define the following indicator function:

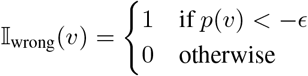

This function assigns a penalty only if the node is confidently predicted to belong to haplotype 2. Homozygous nodes and haplotype 1 predictions are not penalized. (When assembling haplotype 2, we simply negate *p*(*v*).)

The updated beam search scoring function becomes:

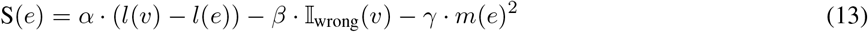

This updated heuristic allows the search to favor paths consistent with grapHiC’s haplotype predictions in the absence of trio-based k-mer information. The exact beam search parameters used in the experiments of this chapter are given in Suppl. Table 6.

## J. Data Availability

As training data, we used the I002 genome published by Sarashetti *et al.* (Sarashetti et al., 2024a) as the basis for simulation. All PacBio HiFi reads were simulated from the reference assembled in (Sarashetti et al., 2024a), while real Omni-C reads were used since these are challenging to simulate adequately in eukaryotic genomes across multiple chromosomes. Raw genomic reads are available in the SRA under Accession No. PRJNA1150503. Further details on the I002 dataset can be found in the associated project repository.

For HG002, both reference and read data are publicly available in the HG002 project repository.

## K. Code Availability

`graphic` and all scripts used for deployment and plotting in this manuscript are available on GitHub at https://github.com/lbcb-sci/grapHiC. The repository also contains the best trained model as reported in the results and a link to our training dataset.

